## Supplementary material for "Trans-Lesion Synthesis and Mismatch Repair Pathway Crosstalk Defines Chemoresistance and Hypermutation Mechanisms in Glioblastoma": NA

#### SUPPLEMENTARY FIGURE LEGENDS

**Supplementary Figure 1. TMZ induced PCNA mono-ubiquitination in nocodazole-synchronized U373 cells**

**(A)** Cell cycle profiles of asynchronous cells, nocodazole-arrested cells, and cells at different times following release from metaphase arrest.
**(B)** Scheme illustrating design of experiment to test effect of TMZ-treatment on PCNA mono-ubiquitination in synchronized cells.
**(C)** Immunoblot showing levels of mono-ubiquitinated PCNA and cyclin E (a late G1/S-phase cell cycle marker) in nocodazole-synchronized cells. 50 μMTMZ was added conditionally to some cultures 11 h following release from the nocodazole block.

**Supplementary Figure 2. Gating strategy for detecting and quantifying mitosin + phospho-histone H3 doubly-stained cells.**

**Supplementary Figure 3. Analysis of *RAD18* expression in patient-derived glioblastomas**

**(A)** Comparison of *RAD18* mRNA expression levels in GBM vs. normal tissue (from TCGA data). Box plots show the lower and upper quartiles. Red: GBM; Gray: normal tissue.
**(B)** Linear regression showing **positive** correlation between *RAD18* expression and proliferation score in GBM (using TCGA data). The Pearson correlation coefficient is shown.
**(C)** Violin plots showing expression levels of RAd18 and other TLS pathway-related genes in malignant glioma.
**(D)** Dot plot showing correlation between *RAD18* expression and proliferation score in recurrent GBM tumors. Samples were stratified according to *RAD18* expression by tertiles. Spearman's rank correlation coefficient is indicated.

**Supplementary Figure 4. Correlation between expression levels of TLS pathway genes and glioma patient survival.** Kaplan-Meier curves showing survival of LGG patients from TCGA stratified by high expression (upper quartile, n=128) and low expression (bottom quartile, n=127) of TLS pathway genes. P values were determined by Log-rank test.

**Supplementary Figure 5. Summary of results from genetic screen**

**(A)** Venn diagram showing sgRNAs targeting DDR genes that were significantly dropped-out (black) or enriched (red) in different genotypes (*RAD18*$^{+/+}$, *RAD18*$^{-/-}$), grown in the presence of TMZ or DMSO (for no-drug control group)
**(B)** Heatmap showing relative enrichment or dropout of sgRNAs targeting MMR genes in different experimental groups.
**(C)** Proposed mechanism for differential dropout of MMR pathway genes in TMZ-treated *RAD18*$^{-/-}$ cells. According to this scheme, *RAD18*-deficiency induces DNA loops that are repaired in a manner involving the MMR loop-resolving complex factors MLH1, MSH3, but not other MMR proteins

**Supplementary Figure 6. Validation of genetic screen for determinants of TMZ-sensitivity in GBM**

**(A)** Experimental workflow of fluorescence-based competitive growth assays

**(B)** Results of competitive growth assays showing TMZ-sensitivities of *FANCD2*, *POLD3* and *PRKDC*- deficient cells.
**(C)** Clonogenic survival assays showing TMZ-sensitivity of WT, *RAD18*$^{-/-}$, *POLD3*$^{-/-}$ and *RAD18*$^{-/-}$ *POLD3*$^{-/-}$ U87 cells.
**(D)** Clonogenic survival assays showing TMZ-sensitivity of WT and *RAD18*$^{-/-}$ U87 cells when grown in the presence or absence of CHK2 inhibitor (ChK2i). TMZ and CHK2i were replenished daily for 5 days. All data points of (C) and (D) represent mean of 3 replicates ± SD; P values were determined by Tukey HSD test.

**Supplementary Figure 7. Summary of sgRNA library design and lentiviral vector used to screen for DDR-dependencies of TMZ-treated GBM cells.**

**Supplementary Figure 8. Workflow of whole-exome sequencing data analysis.**

### Supplementary Figure 1

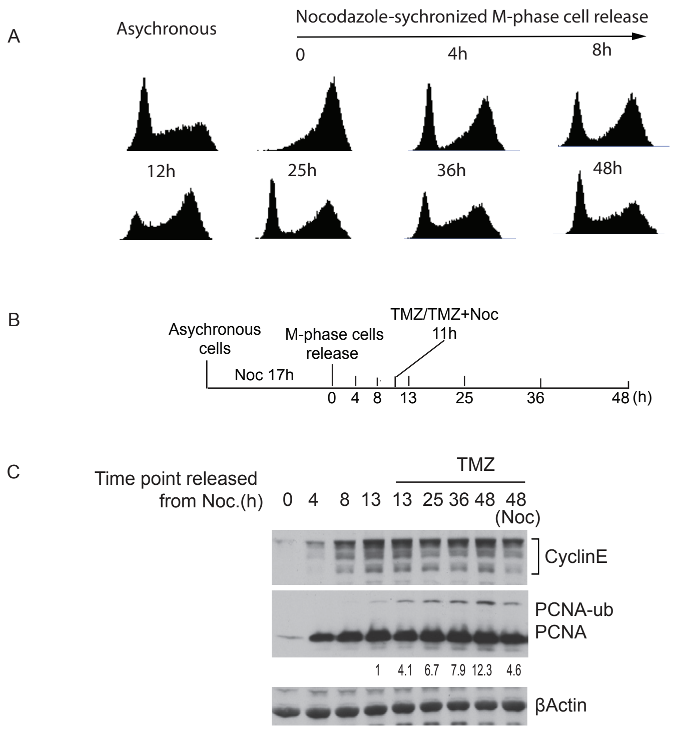

### Supplementary Figure 2

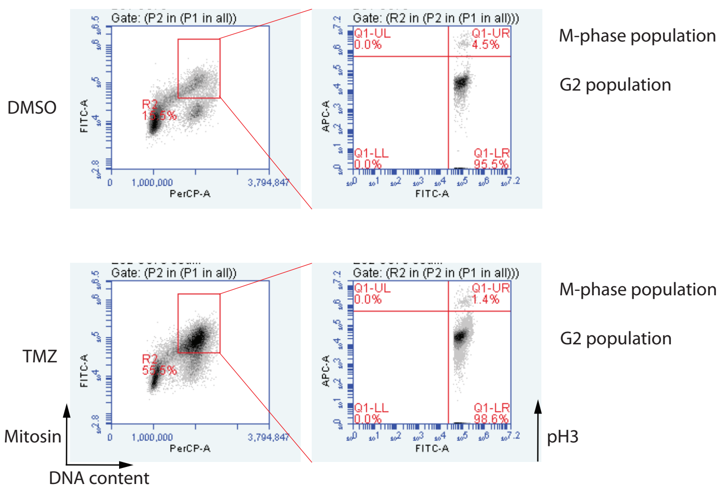

Supplementary Figure 3

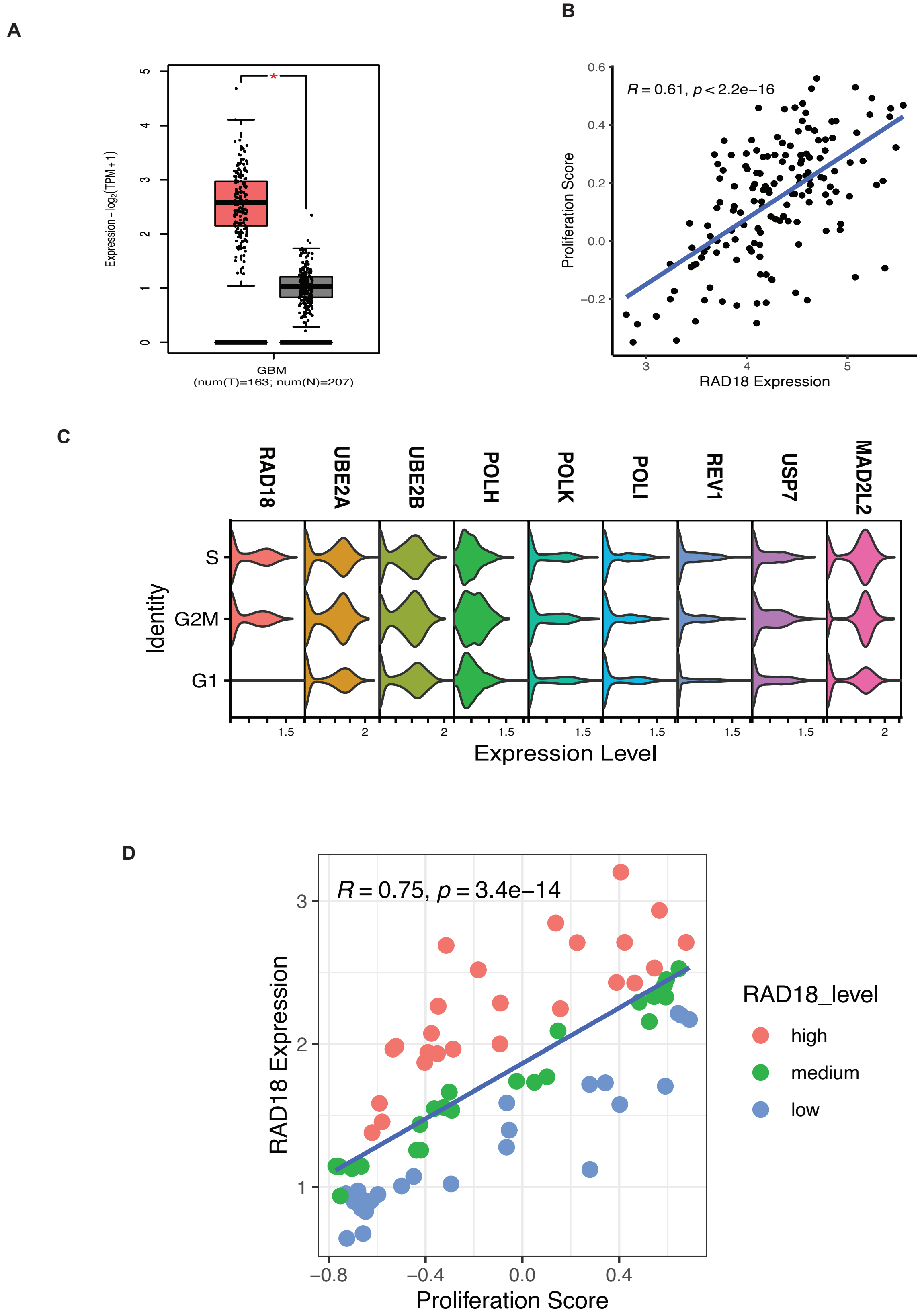

Supplementary Figure 4

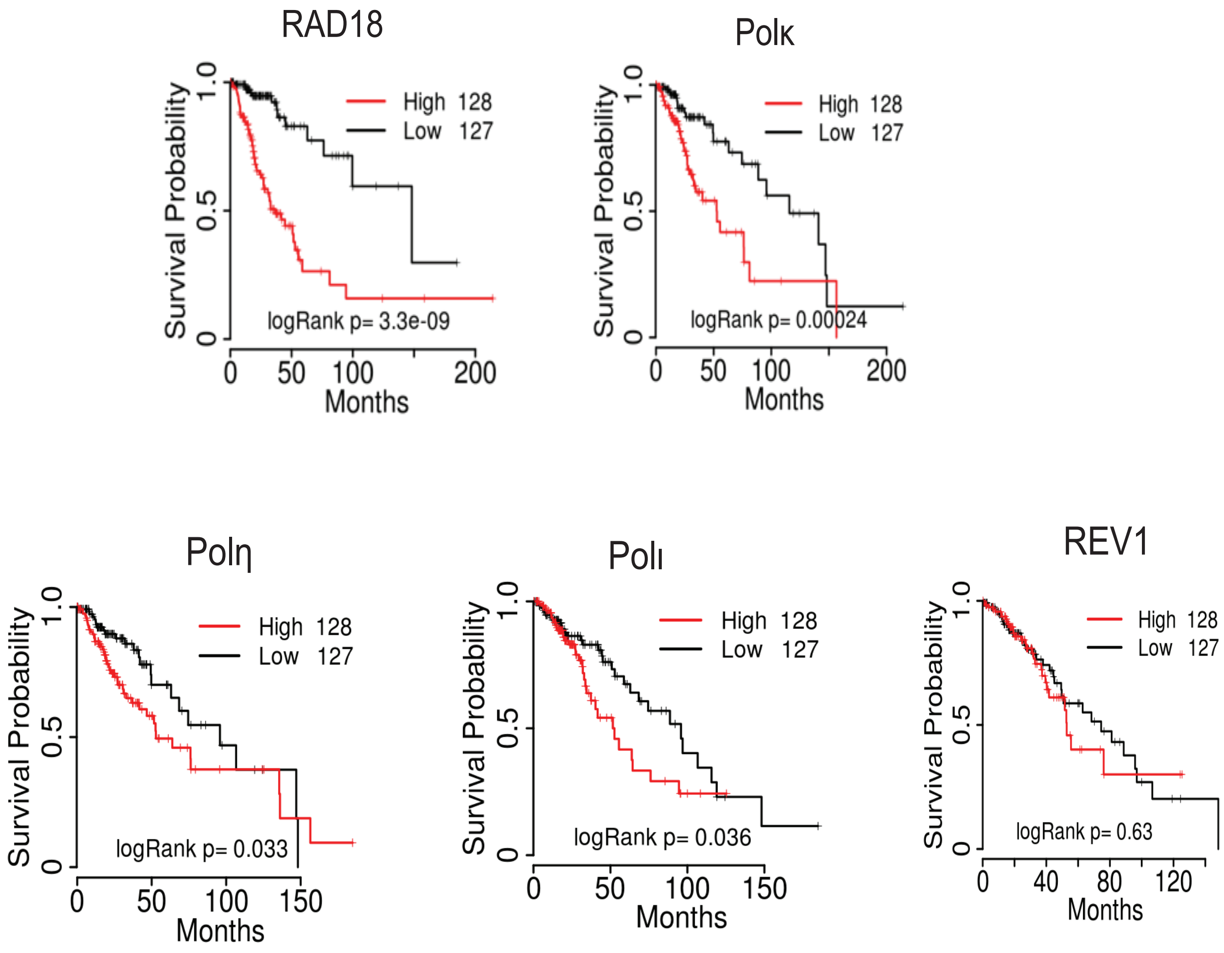

### Supplementary Figure 5

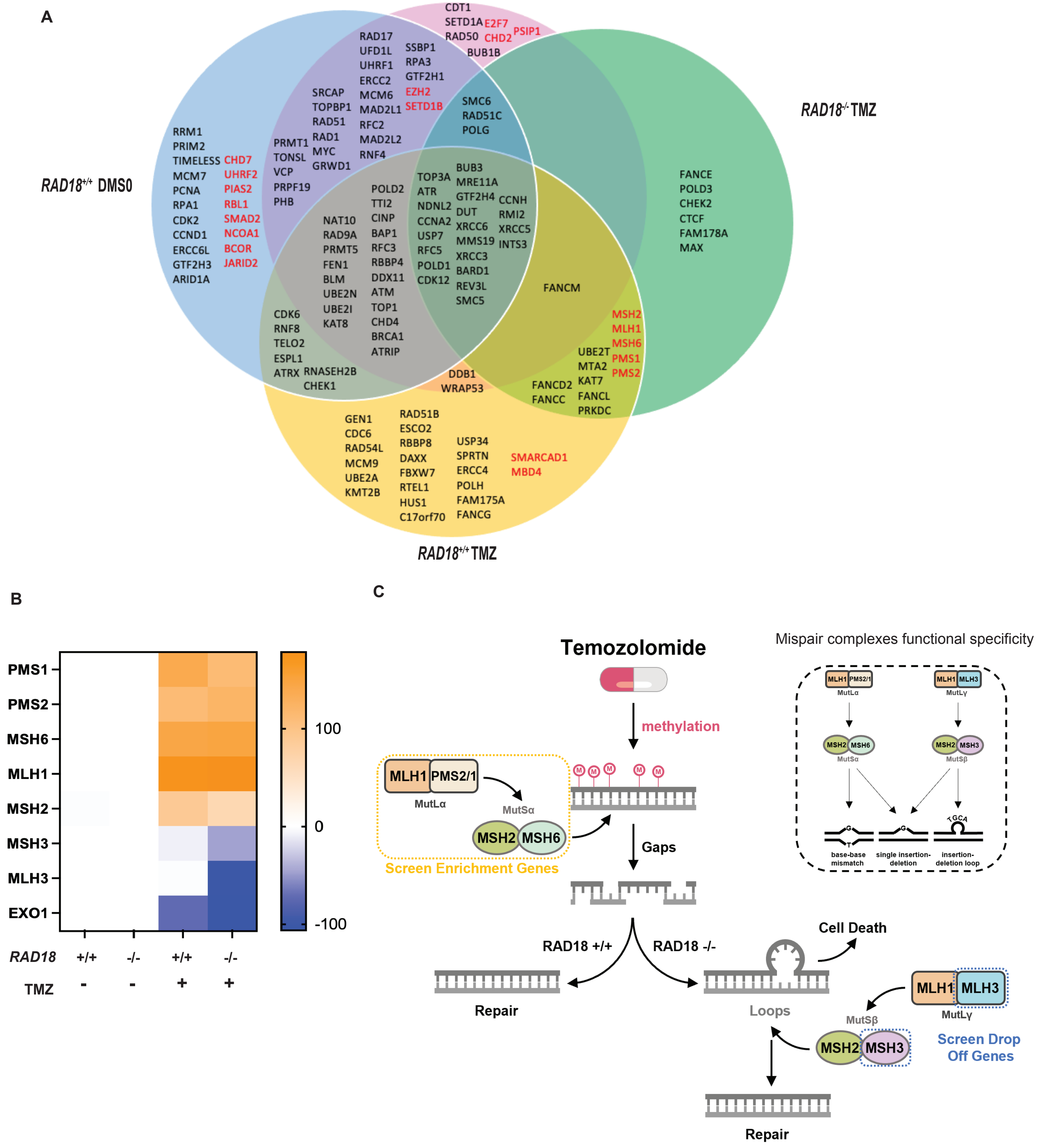

Supplementary Figure 6

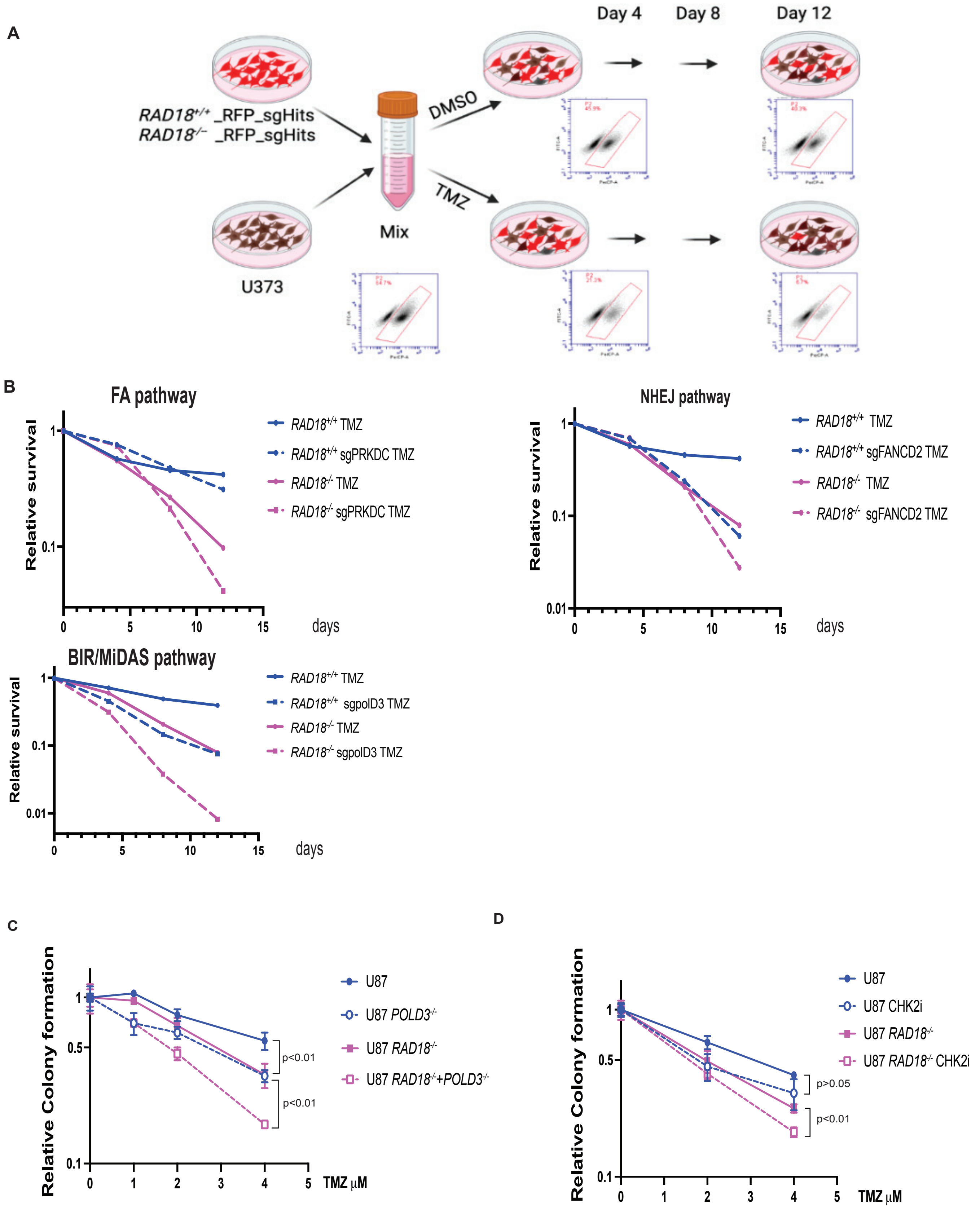

Supplementary Figure 7

1. **Design targeting sites**

Target sequence(20nt) PAM

NNNNNNNNNNNNNNNNNNNNNGG

2. **Oligos to synthesis**

5' Franking sequence | 3' Franking sequence

GTGGAAAGGACGAAACACCGNNNNNNNNNNNNNNNNNNNNGTTTTAGAGCTAGAAATAGCAAGTTAAAATAAGG

3. **Clone oligos to LentiCRISPRv2-DD-Cas9-BSD vector by using 017_ArrayF and 018_ArrayR primers**

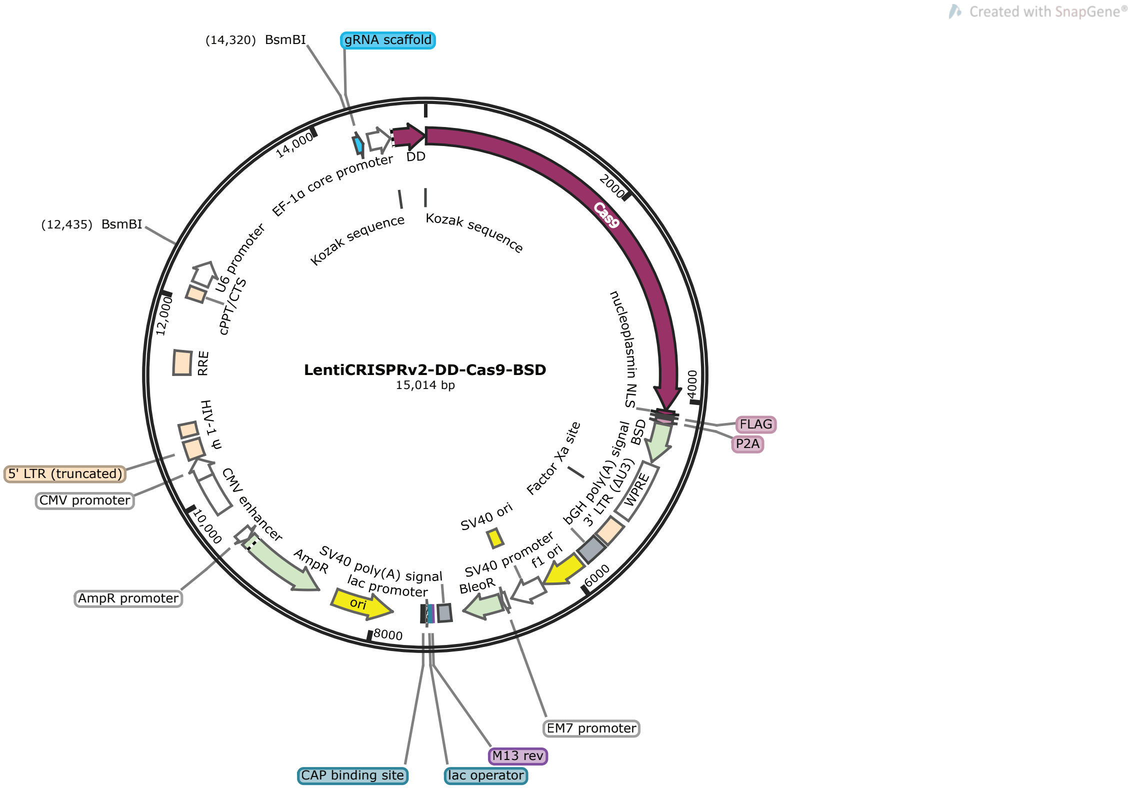


4. **Lentivirus packing**

Supplementary Figure 8

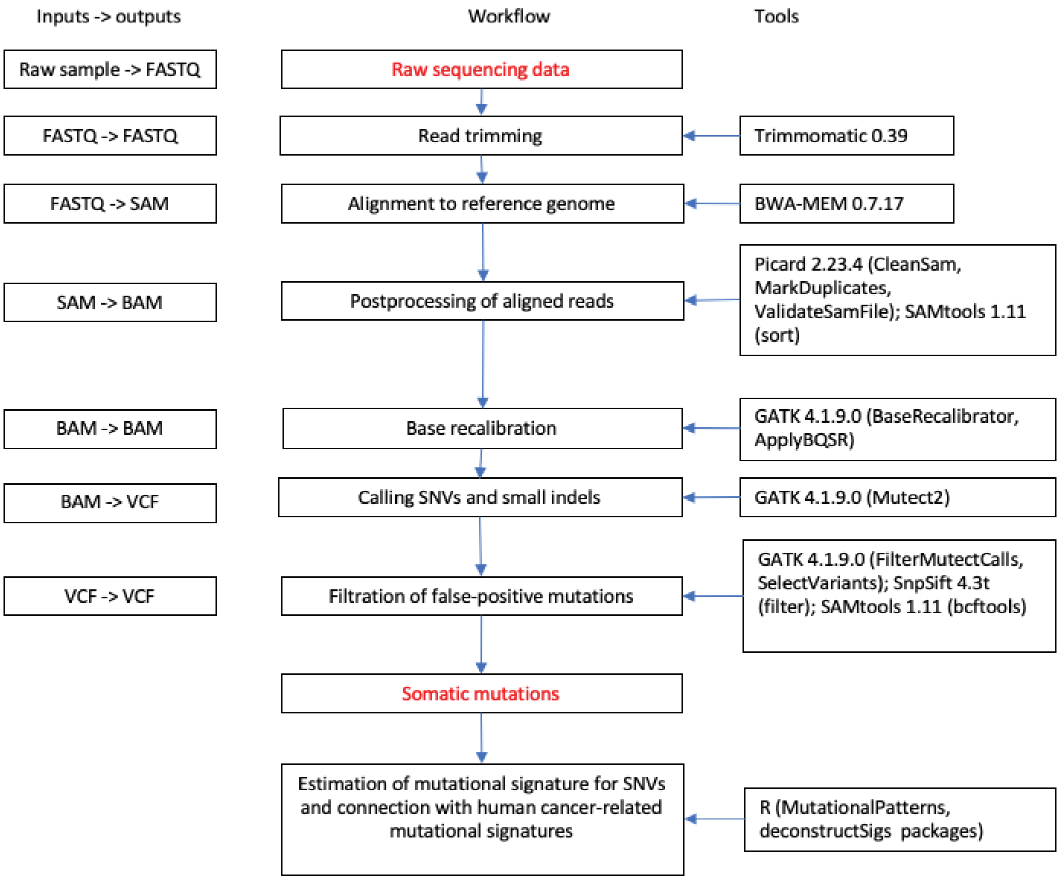
